# The best of two worlds: using stacked generalization for integrating expert range maps in species distribution models

**DOI:** 10.1101/2024.03.12.584552

**Authors:** Julian Oeser, Damaris Zurell, Frieder Mayer, Emrah Çoraman, Nia Toshkova, Stanimira Deleva, Ioseb Natradze, Petr Benda, Astghik Ghazaryan, Sercan Irmak, Nijat Alisafa Hasanov, Gulnar Gahraman Guliyeva, Mariya Gritsina, Tobias Kuemmerle

## Abstract

Species distribution models (SDMs) are powerful tools for assessing suitable habitat across large areas and at fine spatial resolution. Yet, the usefulness of SDMs for mapping species’ realized distributions is often limited, since data biases or missing information on dispersal barriers or biotic interactions hinder them from accurately delineating species’ range limits. One way to overcome this limitation is to integrate SDMs with expert range maps, which provide coarse-scale information on the extent of species’ ranges that is complementary to information offered by SDMs. Here, we propose a new approach for integrating expert range maps in SDMs based on an ensemble method called stacked generalization. Specifically, our approach relies on training a meta-learner regression model using predictions from one or more SDM algorithms alongside the distance of training points to expert-defined ranges as predictor variables. We demonstrate our approach with an occurrence dataset for 49 bat species covering four biodiversity hotspots in the Eastern Mediterranean, Western Asia, and Central Asia. Our approach offers a flexible method to integrate expert range maps with any combination of SDM modeling algorithms, thus facilitating the use of algorithm ensembles. In addition, it provides a novel, data-driven way to account for uncertainty in expert-defined ranges not requiring prior knowledge about their accuracy, which is often lacking. Our approach holds considerable promise for better understanding species distributions, and thus for biogeographical research and conservation planning. In addition, our work highlights the overlooked potential of stacked generalization as an ensemble method in species distribution modeling.

## 1 Introduction

While global biodiversity is declining rapidly (Pimm et al. 2014), our knowledge about species’ distributions often remains limited (Diniz-Filho, De Marco Jr, and Hawkins 2010). This lack of detailed information for many regions and taxa, referred to as the Wallacean shortfall (Hortal et al. 2015), translates not only into knowledge gaps in biogeography and ecology, but also into real barriers for conservation planning to ensure that limited conservation funding is spent most effectively (Hochkirch et al. 2021). Species distribution models (SDMs) have become a central tool for addressing the Wallacean shortfall, allowing to characterize species’ niches by combining occurrence records with environmental predictors for predicting species’ distributions (Elith and Leathwick 2009; Guisan and Thuiller 2005). Yet, although SDMs can accurately assess the environmental suitability of habitats (i.e., map *potential distributions*), they typically lack information on other factors limiting species’ ranges, such as barriers to dispersal or biotic interactions (i.e., competitive exclusion). This, in turn, means that the usefulness of SDMs for mapping *realized distributions* of species can be limited, as their inability to identify range limits often results in an overprediction of species’ ranges, particularly when distributions are modelled across large geographic extents (Calabrese et al. 2014; Merow, Wilson, and Jetz 2017; Soberón 2007). While methods for capturing dispersal and biotic interactions within SDMs have been proposed (Ovaskainen et al. 2016; Zurell 2017), their applicability is often limited due to a lack of adequate datasets or missing knowledge about underlying ecological processes.

A widely-applicable solution to improve SDMs’ capability for assessing realized distributions lies in their combination with external information on species’ range limits (Domisch, Wilson, and Jetz 2016; Fletcher Jr. et al. 2019; Merow et al. 2017). Most commonly, range information is available in the form of expert-based range maps, which offer estimates of species’ range extents derived from occurrence information as well as expert knowledge about geographical, biotic, or environmental range limits. The most important database of range maps (particularly for terrestrial animals) is offered by the International Union for the Conservation of Nature (IUCN), which provides expert-defined ranges for more than 150,000 species (IUCN 2022). Although widely available, expert range maps are frequently criticized for being coarse in resolution (meaning that species will often be absent from many areas within the expert-defined range), incomplete in terms of species coverage or outdated (Higino et al. 2023; Ramesh et al. 2017). Despite these shortcomings, expert range maps often provide the best-available (or only) information on range limits for many species. More importantly, they provide information that is highly complementary to data generated by SDMs (Merow et al. 2017). While range maps characterize a species’ extent of occurrence (i.e., its range limits), SDMs offer fine-scale representations of suitable habitats, making approaches combining both datasets promising for improving distribution assessments (Domisch et al. 2016; Ellis-Soto et al. 2021; Merow et al. 2017).

Several approaches have sought to combine these relative strengths of expert range maps and SDMs, such as using range maps directly as predictors in SDMs (Domisch et al. 2016) or adding spatial offset terms to models that are fit via point process models or related approaches (e.g., Maxent; Merow et al., 2017). The latter approach is particularly promising as it allows to account for uncertainty in expert range maps by incorporating user-defined decay curves that reflect *a priori* expectations about the accuracy of expert range boundaries. Applying this approach, however, can be challenging for two reasons. First, defining spatial offsets and decay curves can be difficult if prior information on the accuracy of range maps is missing, potentially leading to bias introduced by decisions on the strength and decay of the offset term. Second, while the use of algorithm ensembles has become a key approach in species distribution modeling (Araújo et al. 2019; Araújo and New 2007), several widely-used and well-performing machine learning algorithms (e.g., random forests or support vector machines) do not feature offset terms.

Here, we suggest stacked generalization (Wolpert 1992) as an alternative approach for integrating external range information enabling flexible combinations of multiple SDM algorithms. Designed as an ensemble method for combining multiple modeling algorithms, stacked generalization uses the predictions of models built at one level as the input for a meta-learner built at a second level (Naimi and Balzer 2018). Although being widely applied in machine learning (Sesmero, Ledezma, and Sanchis 2015), and despite the general proliferation of algorithm ensembles in SDM studies (Buisson et al. 2010; Hao et al. 2019), stacked generalizations have rarely been used with SDMs (but see Bonannella et al., 2022; El Alaoui & Idri, 2023). Here, we demonstrate the use of stacked generalization as an approach for integrating expert range information with one or more SDM algorithms. Using available occurrence datasets for characterizing expert map accuracy, our approach offers an alternative, data-driven method to integrate expert range maps in SDMs while accounting for their uncertainty.

In the following, we first introduce our approach and highlight issues important to consider in its application. Then, we assess our approach by applying it to a presence-only occurrence dataset for 49 bat species collected across a large geographic extent covering four biodiversity hotspots in the Eastern Mediterranean, Western and Central Asia. Specifically, we compare the predictive performance as well as resulting distribution maps of (1) single-algorithm SDMs, (2) ensembles of SDM algorithms built with stacked generalization, and (3) stacked generalizations including expert range maps.

## 2. Stacked generalization for integrating expert range information in SDMs

Stacked generalization is an ensemble method for combining multiple models, often built with different algorithms, using their individual predictions as training data in a meta-learner (Naimi and Balzer 2018; Wolpert 1992). Here, we apply this approach to integrate one or more SDM algorithms with expert range maps. By using the expert map as an additional model containing complementary information to SDMs (i.e., coarse-scale estimate of range limits), our approach is taking advantage of stacked generalizations working best if heterogeneous input models are combined (Sesmero et al. 2015).

Multiple potential approaches exist to combine SDMs with expert range maps via stacked generalization. One approach is creating a predictor variable for the meta-learner by assigning a fixed value ratio to training points lying inside vs. outside the expert-defined ranges, thereby allowing to control how much weight is given to the expert map (Merow et al. 2017). However, this approach assumes that the probability of occurrence is the same at any distance outside the expert range, although a continuously decreasing probability with increasing distance from the expert range should be expected (Merow et al. 2017). Therefore, we instead use the spatial distance of the training points to the expert range boundaries as a predictor in the meta-learner (Figure 1). This predictor, hereafter referred to as *distance term*, describes the (relative) probability of observing the modeled species within a given distance of the expert-defined range, thereby characterizing the uncertainty of the expert map. This approach is conceptually similar to including spatial offsets with decay curves in point process models (Merow et al. 2017), and results in predicted habitat suitability values smoothly decreasing outside the expert range.

**Figure 1:**
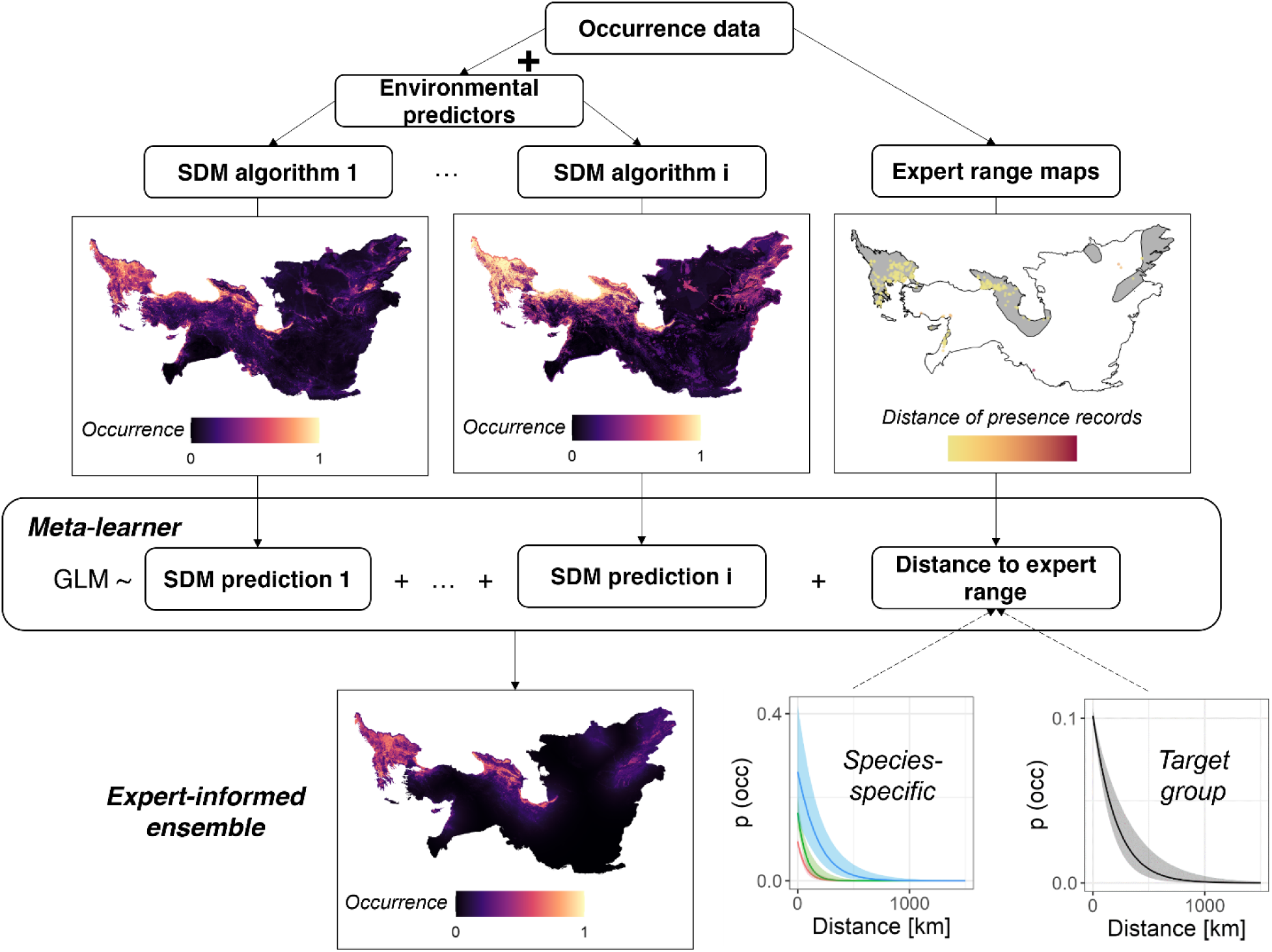
Schematic overview of stacked generalization for combining SDM algorithms with expert range maps. Predictions of multiple SDM algorithms are used together with the distance of occurrence data to the expert range as predictor variables in a logistic regression meta-learner, which then is used to predict the species’ distribution. Maps show examples for one bat species in our dataset (Nyctalus noctula). Map panel for expert range shows IUCN range in grey with presence records colored according to their distance to the IUCN range. Shown maps are in Albers equal area projection.

**Figure 2:**
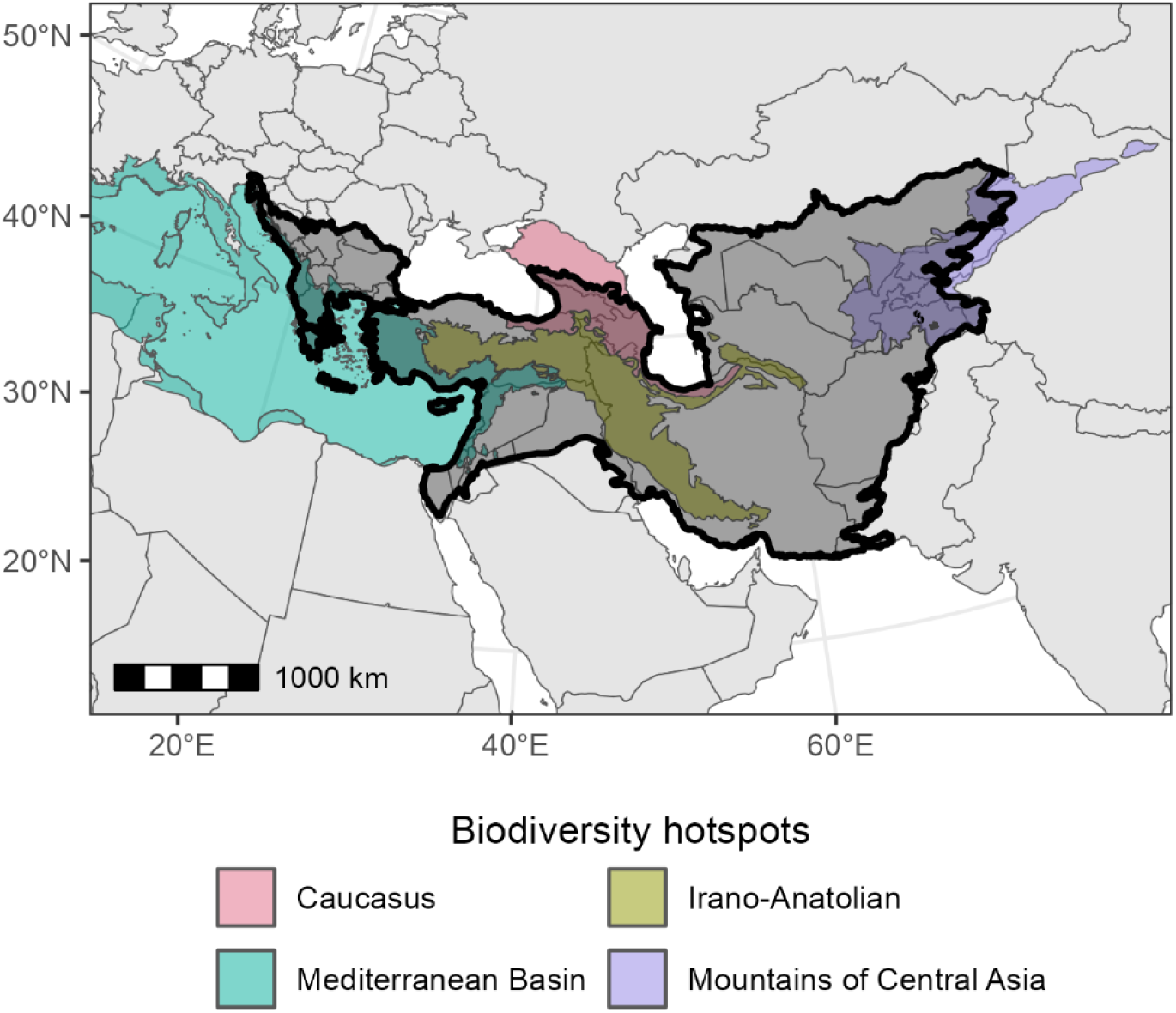
Extent of the study area, shown as black polygon and intersecting global biodiversity hotspots shown as colored polygons. Map is in Albers equal area projection.

However, in contrast to user-defined offsets, the distance term of the meta-learner is derived from the occurrence records used to train SDMs. While using the same datasets for fitting SDMs and assessing the uncertainty of the expert range maps might introduce bias if the collection of occurrence records is influenced by knowledge about expert ranges (Merow et al. 2017), such a data-driven approach will be particularly useful when accurate and representatively sampled occurrence records are available or if prior knowledge about the accuracy of expert range maps is lacking.

While the approach by Merow et al. (2017) allows to control the shape of the decay curve by choosing several curve parameters, in stacked generalizations, the analyst can influence the shape of the fitted distance term through the choice of the meta-learner algorithm or the functional form of the distance term. As a baseline approach we here use logistic regression as a meta-learner, which is widely used in stacked generalizations and results in distance terms following a logistic function similar to the smooth decay curves proposed by Merow et al. (2017). Conceptually, adding the distance term to a logistic regression meta-learner can be seen as adding a constant ‘offset’ to all areas inside the expert-defined range (i.e., areas with distance = 0). This offset is described by the intercept of the logistic regression and expresses the (relative) probability of observing the species inside the expert range given suitability predictions of 0 from all SDM algorithms. Predictions by the meta-learner will decrease with increasing distance from the expert range according to the distance term (Figure 1).

By relating individual species’ occurrences to expert ranges, our approach accommodates species-to-species variability in the uncertainty of expert ranges. However, due to a lack of presence records or highly accurate expert range maps, in some cases only few or no presence records might lie outside expert ranges, which will cause (quasi-)complete separation in the meta-learner. We propose two potential solutions to this issue. First, if species-specific distance terms should be used, bias-reduced logistic regression can be applied for fitting meta-learners (Firth 1993). This commonly recommended strategy for dealing with (quasi-)complete separation in logistic regressions ensures finite parameter estimates and results in responses (i.e., distance terms in our case) that are less steep compared to standard maximum likelihood estimation (Heinze and Schemper 2002). Second, when occurrence data from multiple related taxa are available, species-specific distance terms of meta-learners might be replaced with ‘target-group’ distance terms, which can be calculated by fitting a meta-learner based on training points from multiple or all available species. In this case, the distance term characterizes the uncertainty (probability of occurrences lying outside expert ranges) across all included taxa and does not vary between species, similar to applying the same decay curve across species when integrating range maps as spatial offsets in point process models (Merow et al. 2017).

## 3. Method application

### 3.1 Study area and bat occurrence data

Our study area covers 6.5 million km² and intersects four global biodiversity hotspots (following Myers et al., 2000): The eastern part of the Mediterranean hotspot, the Caucasus hotspot, the Irano-Anatolian hotspot, as well as partially covering the Mountains of Centrals Asia hotspot. To delineate our study area, we fully included all countries in which the sampling of our bat occurrence records was primarily conducted (Afghanistan, Albania, Armenia, Azerbaijan, Bulgaria, Georgia, Greece, Iran, Israel, Montenegro, Syria, Turkey). The borders of our study area were defined based on ecoregion boundaries (Olson et al. 2001). Our study area represents the contact zone between the Western and Eastern Palearctic species pools, where information on the distribution of bats remains limited.

We collected and harmonized bat occurrence datasets from various sources, including national databases, field records, and literature data (see Appendix S1 for an overview of all data sources). In total, we gathered 37,714 occurrence records from 61 taxa. To ensure the quality of records used for model training, we removed all instances in which a species-level identification was impossible or problematic (e.g., uncertain identification within complexes of morphologically highly similar species). In addition, we removed records collected before 1970 to avoid a temporal mismatch between occurrence records and predictor variables (Milanesi, Della Rocca, and Robinson 2020).

Where appropriate, we reclassified records to account for recent genetic analyses that have led to a subdivision of species complexes into multiple cryptic species. This reclassification was done based on available information on the distribution of cryptic species (see Appendix S2 for details on taxonomic revisions within species complexes as well as an overview of species). To remove spatial duplicates and reduce sampling bias, we thinned occurrence records (Boria et al. 2014). As thinning records may reduce model performance for rare species (Steen et al. 2021), we classified species according to the percentile values of sample prevalence (i.e., number of raster cells with presence records) into three classes (low, intermediate, and high prevalence). We then thinned records with minimum distances of 1km, 5km, and 10km for species with low, intermediate, and high prevalence, respectively.

As expert information on species range limits, we compiled IUCN range maps for all species. This led to four species being excluded from modelling since no range map was available. Finally, we selected species with a minimum of 30 remaining records to ensure robust training data sets for building SDMs, resulting in 9,650 presence records from 49 species.

### 3.2 Species distribution modeling

We used presence-background SDMs (Elith and Leathwick 2009) to characterize the distributions of bats in our study area. For modeling, we compiled a set of 40 candidate predictor variables, indicating four key dimensions of habitat suitability for bats: climate, land cover and vegetation productivity, topography and geology, and human pressure and modification (Table 1). While target resolution of our SDMs was 1km², we derived all predictor variables at three spatial scales (1km², 5km² and 10km²), resulting in 120 candidate variables. Including coarser scales derived through moving window averaging allows better characterizing habitat conditions at the scale of bat home ranges (e.g., available forest cover within the surrounding area of a bat roost).

**Table 1:** Overview of environmental predictor variables used in species distribution models.

| Category | Predictor | Available time steps | Data source |
| --- | --- | --- | --- |
| Climate | 19 bioclimatic variables | 1981-2010 (average) | CHELSA climate data (Karger et al. 2017) |
| Land cover and vegetation productivity | Six land-cover proportions (agriculture, forest, shrubs, herbaceous vegetation, bare and sparse vegetation, water) | 1992-2020 (annual) | ESA CCI land cover |
|  | Nine Landsat-based spectral-temporal metrics (cumulative, minimum and seasonality metrics for Tasseled Cap greenness, brightness, and wetness indices) | 1990, 1995, 2000, 2005, 2010, 2015 | Landsat satellite imagery (Oeser et al. 2020) |
| Topography and geology | Terrain ruggedness index | - | (Amatulli et al. 2018) |
|  | Presence of karstifiable rocks | - | World Karst Aquifer Map (Chen et al. 2017) |
| Human pressure and modification | Human modification index | 1990, 2000, 2010, 2015, 2017 | (Theobald et al. 2020) |
|  | Accessibility (travel time to cities) | 2015 | (Weiss et al. 2018) |
|  | Nighttime-lights | 1992-2018 | (Zhao et al. 2022) |
|  | Forest landscape integrity index | 2019 | (Grantham et al. 2020) |

We sampled background points using a target group bias grid, created from kernel density estimation based on all presence records in our dataset. Using the density of bat occurrence records as sampling weights for background points allows characterizing sampling effort and helps to mitigate the influence of sampling bias in presence-background SDMs (Barber et al. 2022; Inman et al. 2021; Syfert, Smith, and Coomes 2013). For each species, we sampled background points equal to ten times the number of available presence records.

We used three SDM algorithms: Maxent (R-package *dismo*; Hijmans et al., 2020), random forests (R-package *randomForest*; Cutler & Wiener, 2022), and boosted generalized additive models (GAMs, R-package *mboost*; Hothorn et al., 2022). Following recommendations by Valavi et al. (2021), we used down-sampled random forests, in which subsamples of the background points are used within each individual tree in order to correct for class imbalances. In a first modeling step, we performed variable selection by fitting univariate models (with default parameters) for all 120 candidate variables and evaluating their predictive performance using the area under the receiver operating characteristic curve (AUC) and Pearson correlation between the predicted and observed presence (COR) in a five-fold cross validation (Valavi, Guillera-Arroita, et al. 2021). For selecting the best-performing model, we combined AUC and COR into a single performance score by rescaling their values across all tested models to a 0-1 scale and calculating the mean of rescaled AUC and COR values. Based on this combined performance score, for each species, we selected the set of variables offering the best predictive performance while having correlation coefficients |r|<0.7 (Dormann et al. 2013). Using the selected variables in a second five-fold cross validation, we tuned algorithm parameters for all species (selecting regularization multipliers for Maxent, *mtry* and *maxnodes* parameters in random forest, and the number of boosting iterations in boosted GAMs; see Appendix S4 for details).

### 3.3 Stacked generalizations

We implemented stacked generalizations in two ways. First, we created pure algorithm ensembles (hereafter *SDM ensembles*) solely relying on the predictions of the three SDM algorithms as predictors in the meta-learner. Second, we created *expert-informed ensembles* additionally including information from IUCN range maps. Additionally, we compared two approaches for adding distance terms to the meta-learner. First, we used species-specific distance terms, using the distance of species-level training points to the species’ IUCN range as a predictor in the meta-learner. Second, we calculated a target-group distance term, which we derived by fitting a logistic regression to the distances of all bat occurrence records in our dataset (i.e., all 49 species). To deal with (quasi-)complete separation in species-specific distance terms, we used bias-reduced logistic regression implemented in the R-package *brglm2* (Kosmidis et al. 2023) for fitting meta-learners.

A critical consideration when using stacked generalizations is the risk of overfitting the meta-learner. A widely adopted strategy for this purpose, referred to as *Super Learner* (van der Laan, Polley, and Hubbard 2007; Naimi and Balzer 2018), uses out-of-sample predictions (i.e., from cross validation) for training the meta-learner. To assess the effect of overfitting on stacked generalizations, we compared meta-learners trained on out-of-sample vs. in-sample predictions (i.e., with and without the *Super Learner* strategy).

We compared the predictive performance of all three tested modeling approaches using five-fold cross validation: (1) single-algorithm SDMs (i.e., Maxent, random forest, and boosted GAMs), (2) SDM ensembles, (3) expert-informed ensembles. To create distribution maps for all species, we predicted all models for the most recent time step (target year for prediction: 2020). To compare mapped distribution patterns between SDM ensembles and expert-informed ensembles, we calculated two metrics: First, species-wise niche breadth using Levins’ B2 metric, describing the uniformity of predicted suitability in geographic space (Warren et al. 2021), and second range overlaps calculated using Schoener’s D metric, describing the similarity of predicted suitability between species pairs (Warren, Glor, and Turelli 2008). We hypothesized that expert-informed ensembles should result in overall lower niche breadths and lower range overlaps, since integrating information on species’ range limits should correct for the overprediction of species ranges by SDMs due to missing information on the effect of dispersal limitations and biotic interactions (Merow et al. 2017).

## 4 Results

The accuracy of IUCN range maps varied considerably across bat species. On average, 73% of presence records fell inside expert-defined ranges (inter-quartile range: 22%), with records lying at an average distance of 50 km of expert-defined range boundaries (inter-quartile range: 30 km). These differences in the accuracy of expert range maps translated into considerable variation in species-specific distance terms and thus clear differences in how predicted suitability values declined outside expert ranges when using expert-informed ensembles. In the case of accurate expert ranges, suitability sharply declined outside expert ranges, leading to the exclusion of (often large) areas identified as environmentally suitable by SDMs but lying outside species’ ranges (e.g., *Myotis myotis* in Figure 3). Conversely, when occurrence records indicated that expert range maps were inaccurate, SDMs clearly dominated the predictions of expert-informed ensembles, allowing to identify areas outside IUCN ranges as likely occupied by species (e.g., *Taphozous nudiventris* in Figure 3). When using target-group instead of species-specific distance terms, suitability values declined at similar rates outside expert ranges across species (Appendix S4).

**Figure 3:**
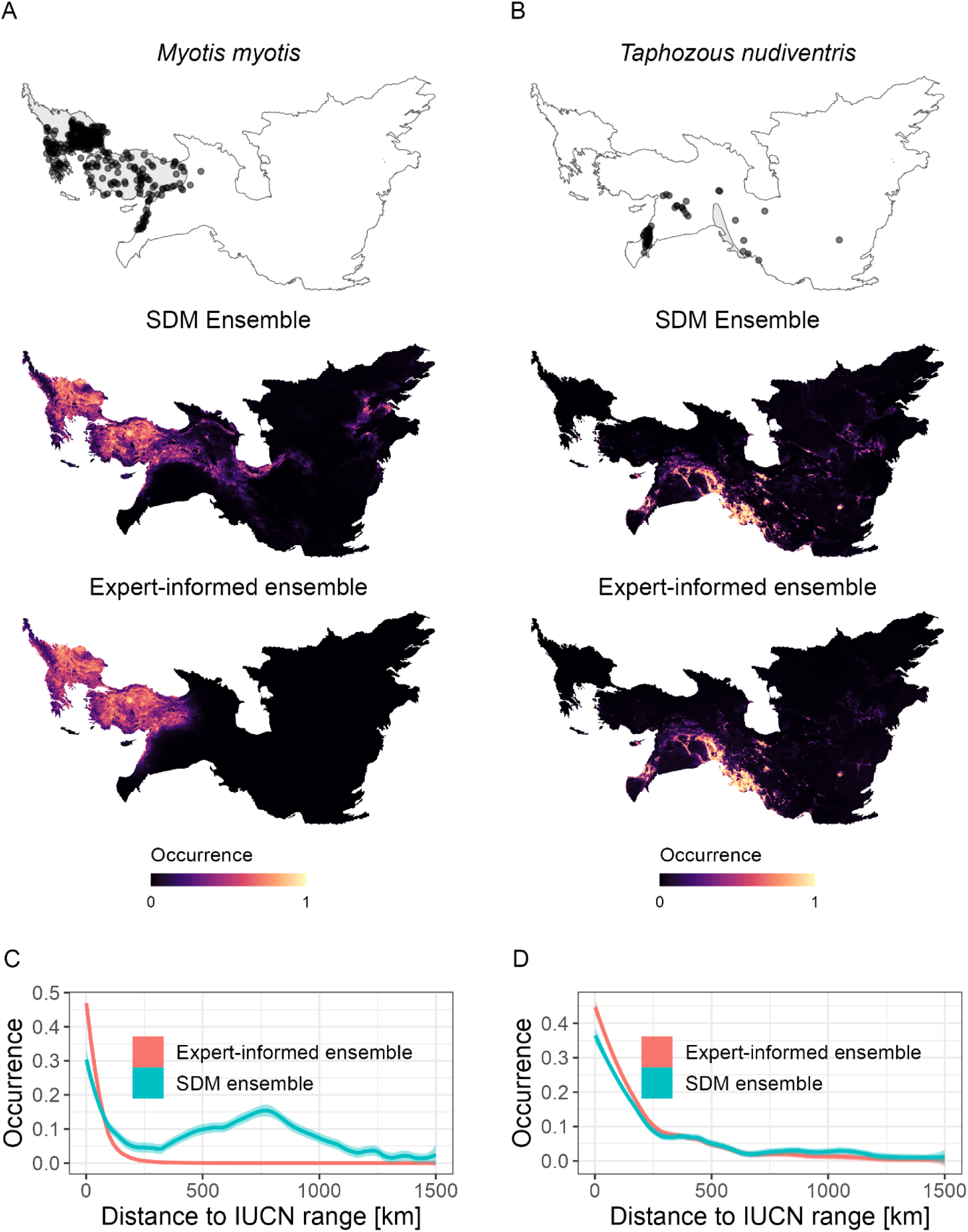
Comparison of distribution maps (A+B) and decline in predicted occurrence probabilities outside expert ranges (C+D) for two example species with high (Myotis myotis) and low expert map accuracy (Taphozous nudiventris). Distribution maps based on IUCN ranges (including available presence records), SDM ensembles, and expert-informed ensembles are shown. Expert-informed ensembles correspond to models built with species-specific distance terms. Plots of decline in predicted occurrence probabilities outside expert ranges (C+D) are based on loess smooth to the data. Maps are in Albers equal area projection.

Considering predictive performance, stacked generalization ensembles outperformed single-algorithm SDMs. However, training on out-of-sample predictions was necessary to achieve optimal predictive performance (i.e., using *Super Learner* approach; Figure 4). Specifically, expert-informed ensembles built with species-specific distance terms achieved the highest predictive performance according to both AUC and COR values, followed by SDM ensembles and expert-informed ensembles with target-group distance terms (Figure 4; since COR values showed no qualitative difference to AUC, we only show AUC values here).

**Figure 4:**
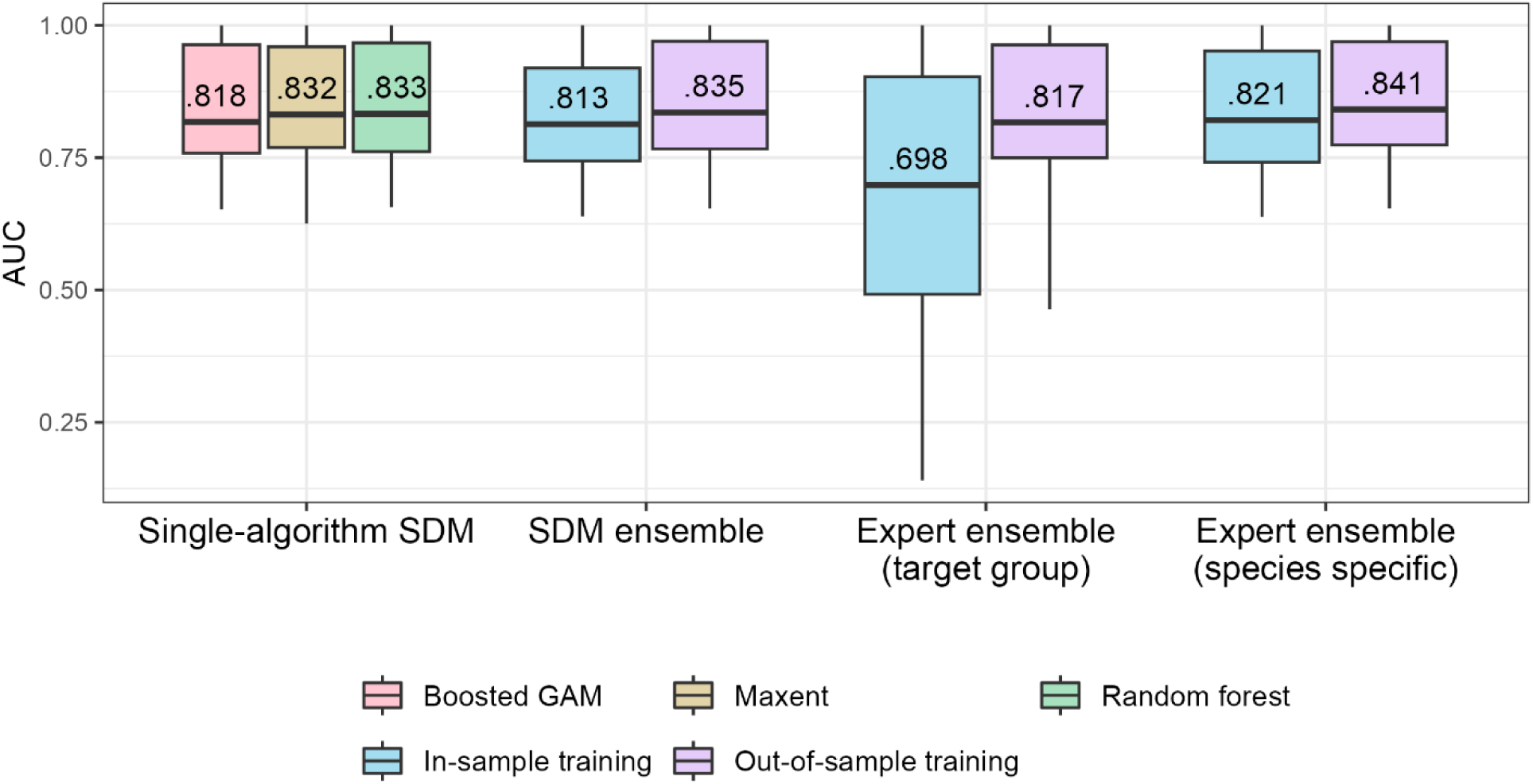
Predictive performance of modeling approaches for 49 bat species in Eastern Mediterranean, Western Asia and Central Asia according to AUC values.

Performance improvements of expert-informed ensembles compared to SDM ensembles generally diminished with the average distance of presence records to expert ranges (i.e., increasing performance gains with higher expert map accuracy; Figure 5A). Performance improvements tended also to be higher for species with fewer available occurrence records as well as for species with smaller range extents (Figure 5B+C), but these relationships were considerably weaker than for expert map accuracy.

**Figure 5:**
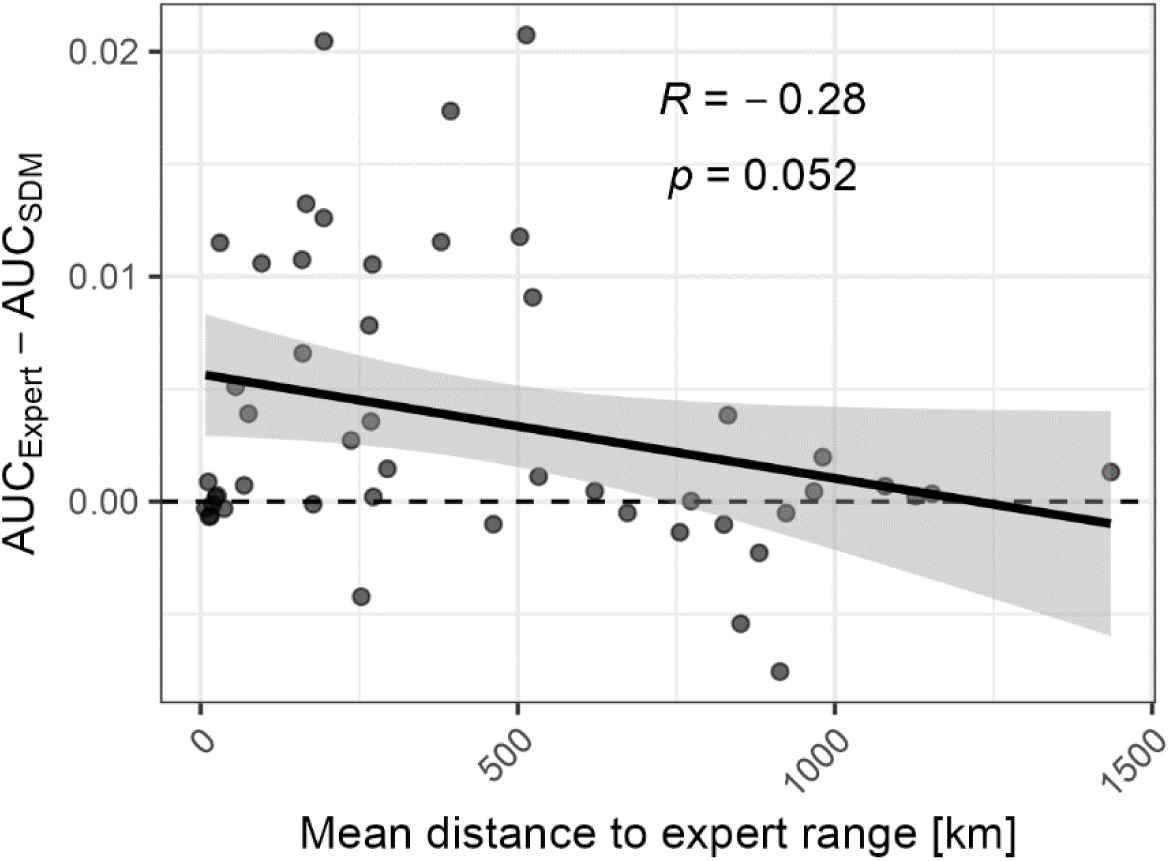
Improvement in predictive performance of expert-informed ensembles compared to SDM ensembles in relationship to expert map accuracy (mean distance of presence records to expert range, including points inside the range with distance = 0). Data for expert-informed ensembles with species-specific distance terms are shown, with linear trend plotted on top.

Considering mapped distribution patterns, expert-informed ensembles resulted in lower niche breadths (i.e., less uniform distribution of predicted suitability in geographic space) for 83% of species compared to SDM ensembles. On average, species-wise niche breadths obtained from expert-informed ensembles were 21% lower compared to niche breadths derived from SDM ensembles (Figure 6A). Range overlaps between species pairs (i.e., similarity of predicted suitability) derived from expert-informed ensembles were lower than overlaps predicted by SDM ensembles in 90% of the cases. On average, overlaps were 30% lower in expert-informed ensembles compared to those obtained from SDM ensembles(Figure 6B).

**Figure 6:**
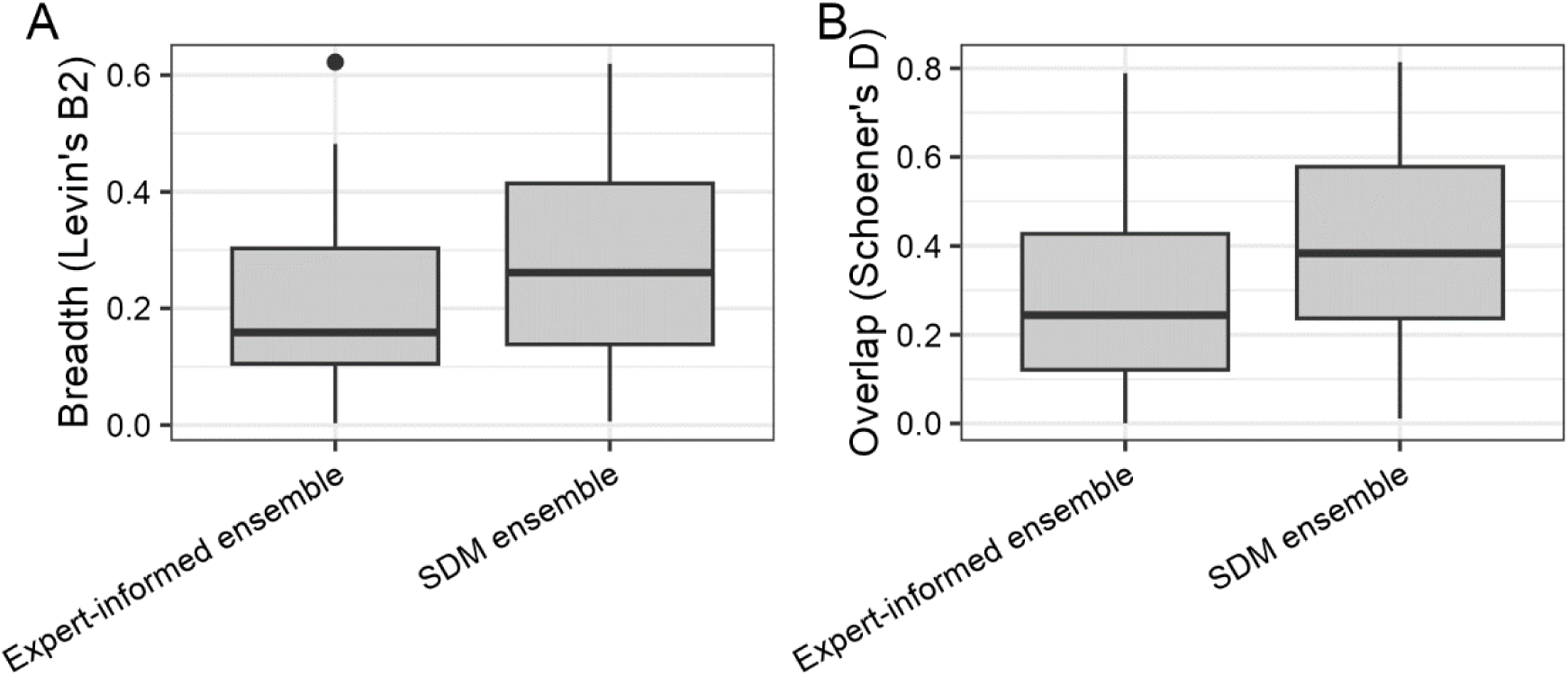
Distribution of (A) niche breadths and (B) range overlaps of bat species according to SDM ensembles vs. expert-informed ensembles.

## 5. Discussion

Addressing the Wallacean shortfall is critical to biogeographical research and conservation planning, yet accurately mapping species’ realized distributions through species distribution modeling presents a significant challenge. Here, we developed a new approach for integrating expert information on range limits in species distribution models by making use of stacked generalization, an ensemble method widely applied in machine learning but still underexplored in the context of SDMs. Testing our approach with a dataset covering 49 bat species demonstrated its flexibility and promise for improving species distribution mapping, allowing to combine the key strength of SDMs (characterizing environmentally suitable habitats) with that of expert range maps (characterizing range limits) without requiring prior knowledge about expert range maps or having to rely on specific modeling algorithms. In a broader context, we add to the growing toolbox of integrated SDM approaches, providing an important step towards more accurate assessments of species’ distributions.

The application of our approach showed that it effectively enables the exclusion of areas lying outside species’ realized range limits, while preserving fine-scale predictions of habitat suitability, which offer a key strength of SDM approaches (Mainali et al. 2020). At the same time, when enough presence records are recorded outside expert-defined ranges, expert range maps exert minimal influence on mapped distributions, demonstrating the flexibility of our approach towards varying levels of expert map accuracy. We did not have an independent validation dataset on species’ absence available, precluding us from performing a more detailed assessment of how our approach affects the accuracy of mapped distributions. Yet, we found improved predictive performance of expert-informed ensembles compared to pure SDM ensembles within our presence-background dataset, with performance improvements depending on the accuracy of expert range maps.

Our approach offers an alternative to applying user-defined spatial offsets in point-process models as proposed by Merow et al. (2017). Choosing between stacked generalization and spatial offsets as ways to integrate expert range maps boils down to selecting different styles of modeling approaches: relying either on prior knowledge when using spatial offsets or on available occurrence datasets when using stacked generalizations for characterizing expert map accuracy. The appropriateness of using stacked generalizations thus hinges on whether available occurrence records can accurately capture expert map accuracy. As highlighted by Merow et al. (2017), using occurrence records for characterizing expert map accuracy requires that their collection is independent of expert range maps (i.e., expert range maps not affecting sampling intensity or species identification), otherwise they will give a biased view on expert map accuracy. However, in many cases occurrence records provide a more comprehensive and up-to-date picture of species distributions compared to expert range maps. Moreover, occurrence records will often be the best available (or only) type of data allowing to evaluate expert range maps, as other *a priori* information on their accuracy is difficult to obtain. Our application of stacked generalizations for 49 bat species highlighted the advantage of allowing for species-to-species variation in the uncertainty of expert-defined ranges, with models achieving the highest predictive performance when using species-specific distance terms. Thus, given that expert map accuracies are expected to vary strongly across species, stacked generalization provides a simple yet effective data-driven approach without the need for manually adjusting prior expectations when assessing many species at once. If occurrence datasets for individual species are deemed too incomplete or biased for characterizing expert map accuracy, target-group distance can be used. Both these options are conceptually very similar to manually defining spatial offsets in point process models based on available evidence on expert map accuracies (Merow et al. 2017), yet eliminate the need for subjective decisions potentially biasing results. In sum, our approach provides an easily and widely applicable data-driven alternative for integrating expert range information in SDMs, proving particularly useful when accurate and comprehensive occurrence datasets are available.

An additional key advantage of our approach lies in its flexibility to combine expert range maps with any combination of modeling algorithms, thereby facilitating the use of algorithm ensembles. In contrast to the use of spatial offsets in point process models, stacked generalizations can be easily combined with machine learning algorithms that do not include offset terms. This enables the use of algorithms such as random forest, often found among the best-performing in comparisons of SDM algorithms (Valavi, Guillera-Arroita, et al. 2021), also featuring the highest discriminative accuracy (i.e., AUC values) in our dataset. With SDM ensembles performing better than any individual modeling algorithm in our dataset, our results also point towards the potential of stacked generalizations as a method for combining modeling algorithms more generally. In line with findings on the importance of avoiding overfitting in stacked generalizations (van der Laan et al. 2007; Naimi and Balzer 2018), we only achieved improved performance when using out-of-sample predictions for training the meta-learner (“Super Learner” approach). It has been shown that in large samples, the Super Learner approach performs at least as well as the best-performing individual algorithm (van der Laan and Dudoit 2003; van der Laan et al. 2007). Yet, despite its potential, stacked generalization has remained neglected in the context of species distribution modeling (El Alaoui and Idri 2023), with studies typically relying on unweighted or weighted model averaging for combining algorithms and stacked generalization not being considered in systematic assessments of SDM ensemble methods (Hao et al. 2020). We therefore recommend stacked generalization as a versatile approach for combining SDM algorithms, which should be included in future comparisons of SDM ensemble methods.

In most cases, the integration of expert range maps resulted in considerably less uniform occurrence predictions and decreased range overlap between species, likely reflecting more realistic predictions of bat distributions in our study area. Both SDMs and expert range maps tend to overpredict occurrence of species since they are missing information on factors limiting species’ ranges (dispersal and competition in the case of SDMs, habitat suitability in the case of expert ranges). Integrating both data sources can therefore improve estimates of individual species’ distributions as well as species richness (Ellis-Soto et al. 2021). Additionally, by disentangling environmental constraints from other limiting factors, the combination of SDMs and expert ranges can help to better understand the influence of non-environmental factors affecting range limits (i.e., biotic interactions and dispersal). For example, contrasting potential range overlaps derived from SDMs with realized range overlaps derived from expert-informed models can provide a window into the potential role of interspecific competition in shaping species’ ranges (Novella-Fernandez et al. 2021). In sum, our approach has broad applicability in ecological research and conservation planning, for example for updating species’ conservation status, assessing conservation priorities through more accurate species richness mapping, and by providing new ecological insights into factors determining species’ range limits.

Our approach adds to the growing toolbox of integrated species distribution modeling approaches by providing a flexible and easily applicable approach for integrating SDMs with readily available information on species’ range limits. As SDMs have become one of the most widely used tools in ecological and biogeographical research, an increasing recognition of their shortcomings has developed (A. Lee-Yaw et al. 2022; Franklin 2010). Recently, integrated modeling approaches have been proposed that try to enhance SDMs by combining them with additional sources of information (Fletcher Jr. et al. 2019; Miller et al. 2019). Integrated SDM approaches already offer key innovations for improving the mapping of species’ realized distributions (Jung 2023; Miller et al. 2019). The broader adoption of these methods, combined with a rapid growth in the availability of biodiversity data will be critical for filling knowledge gaps about the distribution of species and overcome the Wallacean shortfall.

## Acknowledgements

We gratefully acknowledge Alexander Buknikashvili, Christian Dietz, Amit Dolev, Heliana Dundarova, Eran Levin, and Panagiotis Georgiakakis for providing bat occurrence data. Giorgi Sheklashvili, Andrei Kandaurov, Irina Rakhmatulina, Kursantbek Altybaev, Pirimkul Mamatkalykov, Abdurashit Nizamievf, Alexey Dudashvili Svetlana V. Baskakova, Georgiy V. Shakula, Kudaibergen Amirekul, Artemis Kafkaletou Diez, Ioanna Salvarina, and Eleni Papadatou helped with the collection of bat occurrence data. In addition, we are thankful to Christian Dietz for help with the taxonomic classification of bat occurrence records.

