## Appendix for "The best of two worlds: using stacked generalization for integrating expert range maps in species distribution models"

**Appendix S1 Occurrence data sources**

**Table S1**: Number of bat occurrence records by country and data sources used. Data sources are categorized into L = literature data, P = personal field records, and N = national databases. The reference column indicates the publication used for literature data.

| Country | Occurrences | Data sources | Reference |
|---|---|---|---|
| Afghanistan | 92 | L | (Benda & Gaisler, 2015) |
| Albania | 249 | L | (Benda et al., 2019) |
| Armenia | 127 | P | |
| Azerbaijan | 175 | P | |
| Bulgaria | 2114 | N | |
| Cyprus | 185 | L, N | (Benda et al., 2018) |
| Egypt | 65 | P | |
| Georgia | 1007 | N | (Natradze et al., 2023) |
| Greece | 2892 | N | (Georgiakakis et al., 2023) |
| Iran | 623 | L | (Benda et al., 2012) |
| Israel | 569 | N | |
| Jordan | 288 | L | (Benda et al., 2010) |
| Kazakhstan | 22 | P | |
| Kyrgyzstan | 43 | P | |
| Lebanon | 250 | L | (Benda et al., 2016) |
| Montenegro | 160 | L | (Presetnik et al., 2014) |
| Palestine | 156 | N | |
| Romania | 3 | P | |
| Russia | 8 | P | |
| Syria | 172 | L | (Benda et al., 2006) |
| Tajikistan | 8 | P | |
| Turkey | 1396 | P | |
| Turkmenistan | 24 | P | |
| Uzbekistan | 86 | P | |

**Table S2**: Number of bat occurrence records by species.

| Species | Records | Species | Records |
|---|---|---|---|
| Asellia tridens | 83 | Pipistrellus hanaki | 42 |
| Barbastella barbastellus | 155 | Pipistrellus kuhlii | 715 |
| Eptesicus anatolicus | 54 | Pipistrellus nathusii | 175 |
| Eptesicus bottae | 65 | Pipistrellus pipistrellus | 708 |
| Eptesicus ognevi | 37 | Pipistrellus pygmaeus | 231 |
| Eptesicus serotinus | 387 | Plecotus auritus | 130 |
| Hypsugo ariel | 60 | Plecotus austriacus | 149 |
| Hypsugo savii | 456 | Plecotus christii | 63 |
| Miniopterus pallidus | 73 | Plecotus kolombatovici | 70 |
| Miniopterus schreibersii | 434 | Plecotus macrobullaris | 73 |
| Myotis bechsteinii | 136 | Rhinolophus blasii | 282 |
| Myotis blythii | 574 | Rhinolophus clivosus | 52 |
| Myotis brandtii | 64 | Rhinolophus euryale | 302 |
| Myotis capaccinii | 271 | Rhinolophus ferrumequinum | 913 |
| Myotis daubentonii | 137 | Rhinolophus hipposideros | 711 |
| Myotis davidii | 196 | Rhinolophus mehelyi | 101 |
| Myotis emarginatus | 370 | Rhinopoma cystops | 87 |
| Myotis myotis | 333 | Rhinopoma microphyllum | 70 |
| Myotis nattereri | 179 | Rhinopoma muscatellum | 33 |
| Nyctalus lasiopterus | 86 | Rousettus aegyptiacus | 167 |
| Nyctalus leisleri | 218 | Tadarida teniotis | 417 |
| Nyctalus noctula | 253 | Taphozous nudiventris | 52 |
| Nycteris thebaica | 29 | Taphozous perforatus | 35 |
| Otonycteris hemprichii | 74 | Vansonia rueppellii | 27 |
| | | Vespertilio murinus | 123 |

**Appendix S2 Taxonomic revisions of occurrence records**

In five species groups, we updated the species identifications to reflect taxonomical changes.

1) The whiskered bats group, *Myotis mystacinus* sensu lato:
   a. We reclassified *M. aurascens* as *M. davidii* (Benda et al., 2012), and further included *M. hajastanicus* into *M. davidii* (Dietz et al., 2016).
   b. Based on the genetic analysis, *Myotis mystacinus* appears to be relatively rare in the study area (Çoraman et al., 2020). Additionally, all the analyzed *M. davidii* individuals from this region carry *M. mystacinus* mitochondrial DNA, which further complicates the species identification based on mitochondrial markers. Therefore, to avoid misidentifications, we only used *M. mystacinus* records which were genetically confirmed (Çoraman et al., 2020) or were identified based on skull measurements (Benda et al., 2016; Benda & Karataş, 2005)
2) We adopted the revised species identifications of *Myotis nattereri* sensu lato as *M. nattereri*, *M. hoveli* and *M. tschuliensis*, following (Çoraman et al., 2019), (Kruskop & Solovyeva, 2021), and (Uvizl & Benda, 2021). Samples lacking genetic confirmation from the putative contact zones were excluded.
3) *Miniopterus schreibersii* records from the eastern distribution ranges were revised as *M. pallidus* following (Furman et al., 2009) and (Šrámek et al., 2013). Samples lacking genetic confirmation from the putative contact zones were excluded.
4) In the *Plecotus* group, we revised the identification of specimens previously assigned to *P. auritus* and *P. austriacus*, as some records predated the descriptions of *P. kolombatovici* and *P. macrobullaris*.
5) In the *Eptesicus* group:
   a. Anatolian records of *E. bottae* are regarded as *E. anatolicus*.
   b. *Eptesicus turcomanus* was in included in *E. serotinus* following (Juste et al., 2013).

**Appendix S3 Tuning of hyperparameters in SDM algorithms**

**Table S4**: Hyperparameter settings tested for tuning SDM algorithms. In random forest models, all combinations of both tuning parameters were tested, resulting in 20 tested settings per algorithm.

| | **Maxent** | **Random forest** | **Boosted GAMs** |
|---|---|---|---|
| **Parameter** | Regularization multiplier ($\beta$) | Maximum tree depth (*maxnodes*),<br>Number of variables randomly selected at each split (*mtry*) | Number of boosting iterations (*mstop*) |
| **Tested settings** | 0.5, 1, 1.5, 2, 2.5, 3, 3.5, 4, 4.5, 5, 6, 7, 8, 9, 10, 12, 14, 16, 18, 20 | 5, 10, 15, 20 (maxnodes)<br>2,4,6,8,10 (*mtry*) | 40, 50, 60, 70, 80, 90, 100, 150, 200, 250, 300, 350, 400, 450, 500, 600, 700, 800, 900, 1000 |

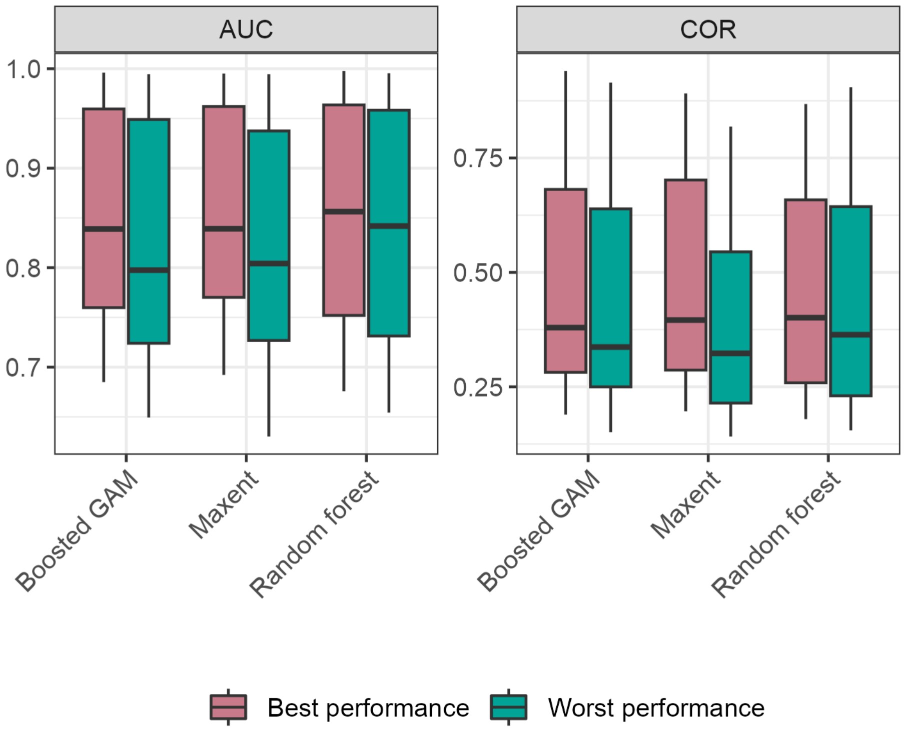


***Figure S1***: *Effect of hyperparameter tuning on performance of SDM algorithms. Plot shows predictive performance of the three SDM algorithms across 47 modeled bat species according to the AUC and COR metrics, comparing the selected, best-performing hyperparameter setting with the worst-performing one.*

### Appendix S4 Distribution maps obtained with target-group distance terms

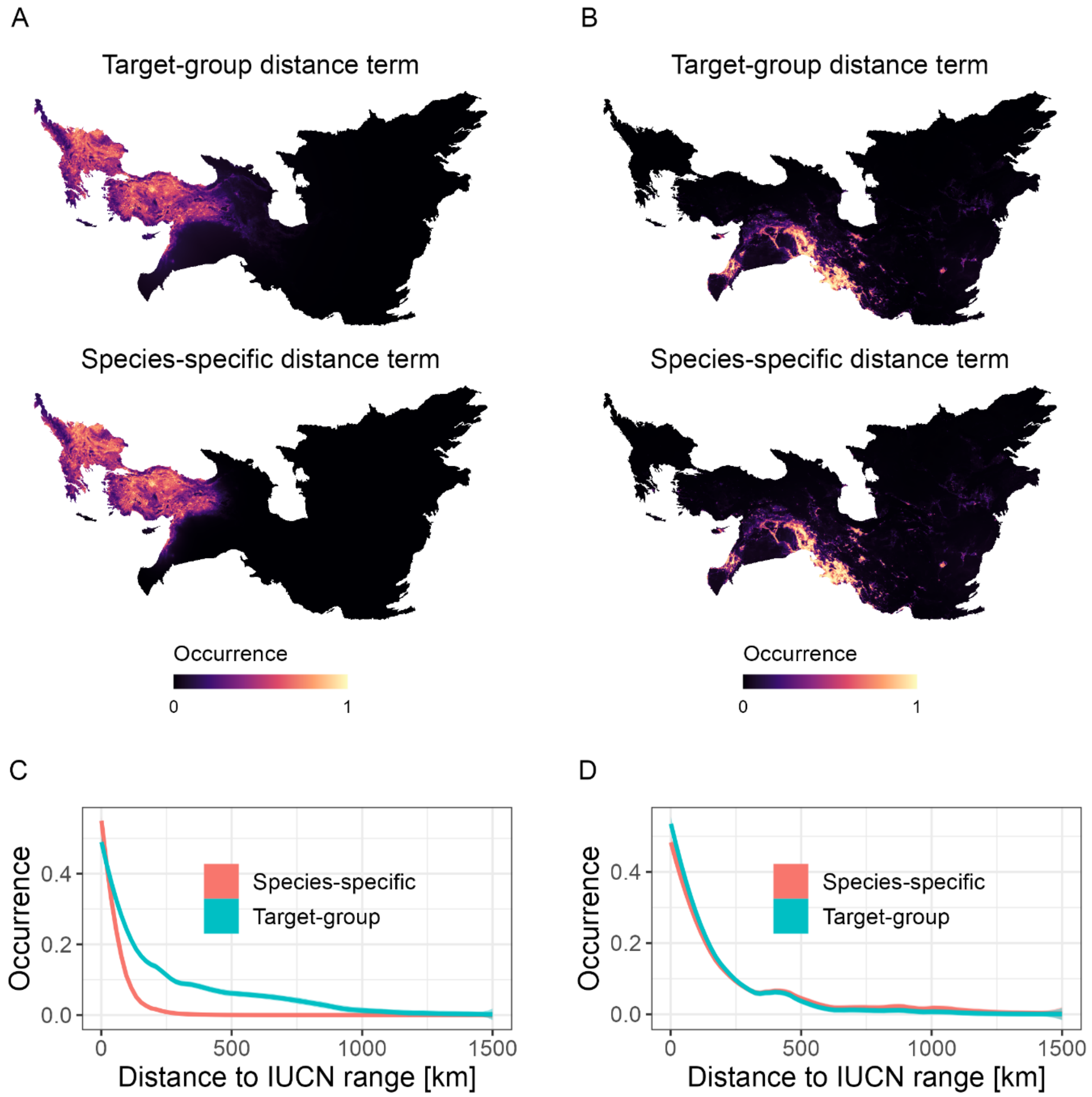


**Figure S2**: *Comparison of distribution maps obtained from different expert-informed ensembles (A+B) and decline in predicted occurrence probabilities outside expert ranges (C+D) for two example species with high (Myotis myotis, left column) and low expert map accuracy (Taphozous nudiventris, right column). Distribution maps show predictions of expert-informed ensembles built with target-group and specific-specific terms, respectively. Plots of decline in predicted*

*occurrence probabilities outside expert ranges (C+D) are based on loess smooth to the data. Maps are in Albers equal area projection.*
